# Functional niche attributes shape species interaction roles in ecological networks

**DOI:** 10.64898/2026.09.13.751296

**Authors:** Georg Albert, Andrea Davrinche, Helge Bruelheide, Felix Fornoff, Sylvia Haider, Werner Härdtle, Lin Jiang, Alexandra-Maria Klein, Yi Li, Xiaojuan Liu, Keping Ma, Goddert von Oheimb, Tobias Proß, Michael Staab, Ming-Qiang Wang, Pandeng Wang, Xian Yang, Chao-Dong Zhu, Andreas Schuldt

## Abstract

Identifying how species interact in ecological networks is key to understanding species’ contributions to ecosystem functioning, but can be difficult to establish in species-rich communities. Functional niches, which capture broader ecological strategies, are often easier to assess and could help approximate species’ interaction roles. We tested this by integrating species interaction and leaf functional trait data of tree species and leaf-associated interaction types from a large-scale forest biodiversity experiment. We found that identity, variability, and similarities of functional niches shaped corresponding aspects of interaction roles. Interestingly, however, this association inversed in the case of functional niche variability, which decreased instead of increased the variability of species interactions. This suggests a potential trade-off between processes that determine variability in functional and interaction space. Furthermore, effects were not constrained within aspects, with similarities of interactions being particularly responsive to most tested niche aspects, indicating that interaction-based competition among plants is particularly sensitive to functional niche differences. Moreover, many effects of functional niches depended on tree species biomass and tree species richness, highlighting the importance of the larger ecological context for determining species interaction roles. Together, our findings show that species’ interaction roles, while clearly altered by, are not simply a consequence of functional niches. Instead, interaction roles integrate multiple aspects of functional niches depending on species performances and community composition and can therefore be crucial for anticipating ecosystem change.

## Introduction

Species interactions are an integral part of ecosystems. Interactions determine the allocation and transfer of energy, nutrients, and biomass across trophic levels (e.g. plants, herbivores), thereby defining ecosystem functions (e.g. plant productivity) and associated services (e.g. carbon storage; Albert *et al*. 2026; Barnes *et al*. 2018). The way a species interacts with other species thus determines its impact on ecosystems, rendering species’ roles in interaction networks (i.e. how they interact with other species) critical for assessing their contributions to ecosystems and anticipating ecosystem change (Cirtwill *et al*. 2018). For example, specialized species that interact with few other species and species with more ubiquitous interaction partners can simultaneously enhance the efficiency of ecosystem processes and completeness of resources used, thus support multiple ecosystem functions (Albert *et al*. 2026). However, explicitly sampling all interactions to identify species interaction roles is an often labour- and time-intensive endeavour. Since interaction roles are inherently linked to functional niches (Cirtwill *et al*. 2018; Coux *et al*. 2016), approximating them from species’ functional niches is an attractive alternative approach.

Usually estimated using functional traits, functional niches can align with interaction roles by integrating traits that determine interactions (e.g. mandibular traits of insects; Ibanez *et al*. 2013). Accordingly, functional traits have seen wide application for deriving species interactions as they reflect ecological strategies of species. For example, many predator-prey interactions include predators that are larger than their prey (Brose *et al*. 2019), making body size an important predictor of predator-prey interactions. Successful pollination can similarly depend on matching pollinator and flower traits, such as beak length and corolla depth in pollinating birds (Peralta *et al*. 2024). Such trait-matching approaches have a strong mechanistic underpinning, utilizing traits that are often specific to interaction types. Species, however, usually interact with a diverse set of functional groups, with interaction partners ranging from microbes to mutualists, parasites, and other higher-level consumers. For each group, functional traits fundamentally define which interactions could be realized. Instead of focusing on trait-explicit associations for each group, identifying how species interaction roles result from functional niches offers a shortcut that allows generalizations across interaction types. However, functional niches usually capture broader ecological strategies based on a wider set of functional traits that may only indirectly affect species’ interaction roles. The explicit identification of species’ interaction roles from species functional niches can therefore be challenging.

While functional niches are measured in functional trait space (Violle & Jiang 2009), the interaction role of species is derived from networks of species interactions (Cirtwill *et al*. 2018). Despite this difference, similar aspects are quantifiable. Identity and variability of species’ functional traits are commonly used to describe the relative position and breadth of functional niches (Violle & Jiang 2009). Positioning species along trait-based gradients is a particularly widespread approach to define a species’ functional identity, with the most prominent example being the leaf-economic spectrum that separates acquisitive and conservative growth strategies of plant species (Wright *et al*. 2004). Compared to acquisitive species, plants with conservative growth strategies tend to invest more into structural elements and secondary metabolites. These can, for example, act as defence mechanisms against herbivory (Coley *et al*. 1985), or determine the suitability of a plant as a habitat for mutualistic species such as endophytic bacteria (Tellez *et al*. 2022). Such trait-based shifts in a species’ niche position can directly translate into shifts of its interaction role. For example, a shift of the functional identity of a species could also shift its identity in an interaction network. The identity of a species in an interaction network can be defined as the proximity to other species (i.e. closeness) and by assessing how much a species connects different species in the network (i.e. betweenness). These measures characterise the identity of a species as its centrality in a network. Species with a high centrality are key mediators of perturbations from biotic and abiotic stressors, determining ecosystem stability directly and through higher order interactions (Martins *et al*. 2024, Albert *et al*. in prep). Interestingly, the degree of a species, which captures network centrality as the number of interaction partners, describes the variability of a species’ interactions and closely relates to a species’ identity in the network (Jordán *et al*. 2007). This suggests that centrality metrics capturing a species’ identity and variability in interaction networks may show similar responses to functional niches.

Aspects of species’ functional niches may not exclusively affect similar aspects of species’ interaction roles. This becomes particularly apparent when considering not only identity and variability of species but also similarities between species. Similarities between species’ functional niches are often interpreted as a sign of competition (e.g. Mason *et al*. 2011). Species that share interaction partners with other species compete via their shared interactions (Holt 1977). However, while similarities of interactions between species may follow similarities in functional niches (low functional uniqueness), interactions can also be more similar for species with average functional strategies (low functional specialization) and broader functional niches, hence be influenced by functional identity and variability, respectively. Effects of multiple and not just matching aspects of functional niches on interaction roles would therefore indeed be expected. Furthermore, effects of functional niches on interaction roles do not need to be strictly coordinated and could also display trade-offs or neutral relationships. For example, if a plant community is structured by limiting resources, plant species can avoid competition by differentiating functional niches (Tilman *et al*. 1997). Unless species interactions are similarly limiting, they can remain largely unaffected by such shifts of the functional niche, and may even display inverse trends as they help avoid competitive exclusion (Albert *et al*. 2022; Brose 2008). Hence, effects of species’ functional niches on interaction roles are not necessarily constrained to the same aspects (identity, variability, similarity), and may additionally reflect trade-offs between processes that limit coordinated associations.

How functional niches and interaction roles are coordinated, displaying trade-offs or varying synchronously, depends on the larger community context. Biodiversity, and species richness in particular, fundamentally defines which species can interact, modifying competition and the need to differentiate niches (Albert *et al*. 2026; Blüthgen & Staab 2024). Especially when species richness is high, species with more unique functional strategies have a competitive advantage (Mahaut *et al*. 2023). A high species richness simultaneously favours generalist interaction partners that can spill-over from other species (Borremans *et al*. 2019). Functionally unique species that have limited interaction partners in low richness communities could therefore end up playing a very similar interaction role as less functionally unique species at high species richness.

Due to their position at the foundation of many interaction networks, as well as their relatively large contribution to biomass, especially in terrestrial systems, plant species occupy a very unique position in interaction networks (Bar-On *et al*. 2018). Because more productive plant species provide more resources to their interaction partners, the role a plant species plays in interaction networks is not only defined by its functional niche but also by its biomass. In particular, plant biomass modifies the effects of functional niches on interaction roles. For example, productive species likely occupy functional niches that are associated with more acquisitive growth strategies (Bongers *et al*. 2021). Alternatively, species may be more productive because they minimize impacts from unfavourable environmental conditions, such as pests, which is associated with conservative growth strategies (Augusto *et al*. 2025). The interaction partners associated with either plant strategy can differ drastically, potentially shifting the interaction roles of plant species (Neyret *et al*. 2024). In contrast, less productive species that have an overall smaller capacity to interact may also vary less in their interaction role regardless of their growth strategy. For plants in particular, the effect of functional niches on interaction roles can therefore differ depending on the biomass, capturing shifting trade-offs or synchronous variation between them.

To investigate how functional niches of plant species determine plant interaction roles, we synthesize trait and interaction data collected for 19 tree species planted in a large-scale tree biodiversity experiment in subtropical China (BEF-China; Bruelheide *et al*. 2014). To gain generality, we not only use a diverse set of tree species but also integrate three types of interactions (Lepidoptera herbivores, leaf-sucking Hemiptera, endophytic bacteria; Fornoff *et al*. 2019; Wang *et al*. 2019; Yang *et al*. 2023), all relying on tree leaves as resource and habitat. To match this, we characterized plant functional niches using leaf functional traits, explicitly incorporating within-species differences. Specifically, we use indices that capture three different aspects of functional niches: species identities (positions along principal component axes, and distance to centre of trait space), species variation (size and evenness of convex hull spanned up by a species in trait space), and species similarities (uniqueness of occupied functional trait space; Fig. 1a-b). We furthermore captured plants species’ interaction roles using measures that similarly capture species’ identities (closeness and betweenness), variation (degree and evenness of interactions), and similarities (similarities of interactions of the focal plant species and their consumers, i.e. consumer dependence; Fig. 1c). We then test if the interaction roles of the 19 tree species can be explained by their functional niches, considering interactions with tree species richness, i.e. the experimental treatment of BEF-China, and tree species biomass. We expect that (1) indices related to specific aspects of tree species’ interaction roles (i.e. identity, variability, similarity) are best explained by indices capturing the respective aspects of the functional niche. However, we further expect that (2) interaction roles can additionally respond to multiple aspects of functional niches. Finally, we expect that (3) tree species richness and tree biomass modify the association between functional niches and interaction roles, with the former playing a particularly pronounced role for similarity-based metrics, and the latter potentially influencing a wider range of associations.

**Fig. 1:**
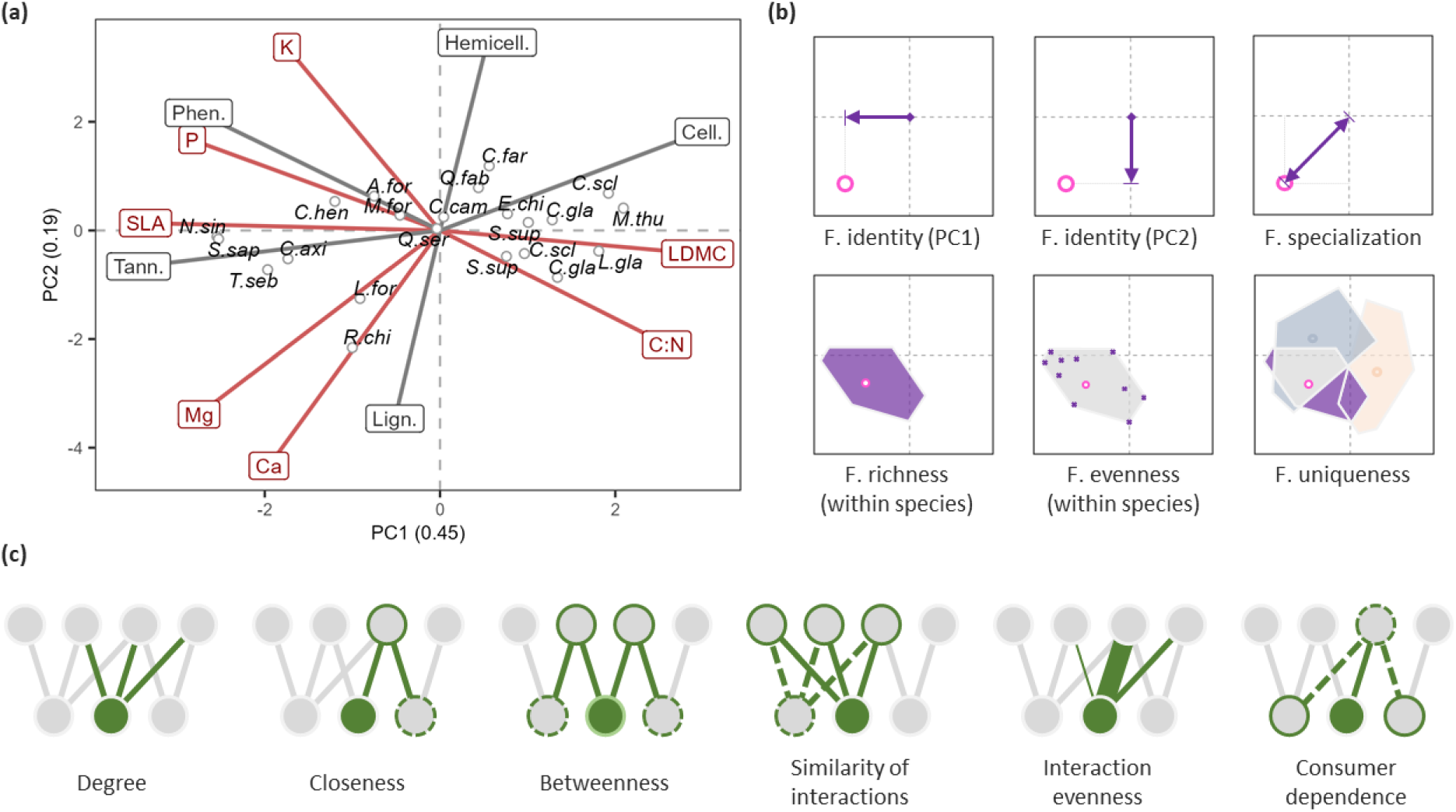
Metrics capturing functional niches, extracted from functional trait space, and interaction roles of tree species. (a) Principal component analysis (PCA) based on functional traits of the included species. Species position in ordination space is indicated as circles, with abbreviations of species names (see Tab. S1 for full names). Functional traits are shown as lines, with red traits being used for calculating the PCA, and grey traits fitted post hoc (see Methods). (b) Illustration of functional niche metrics for a focal species (pink circle). Functional identity is defined as the position on the two principal component axes, indicated as arrows. Functional specialization is measured as the distance to the centre of the trait space, thus capturing how much a species deviates from an average species. Polygon illustrates within-species variability used to measure functional richness. Functional evenness captures how evenly values are distributed in the convex hull of a species. Functional uniqueness measures dissimilarities between species, i.e. the non-overlap between convex hulls. (c) Illustration of interaction role metrics. Focal species is shown as filled green circle. Degree captures the number of interactions of a species. Closeness and betweenness respectively quantify the distance of species to other species, and how often a focal species lies on the shortest path between species pairs. Similarity of interactions is the average overlap of interaction partners. Interaction evenness is Shannon evenness of interaction strength of a species. Consumer dependence is the relative importance of a species for its consumers. A more detailed description of the functional niche and interaction role metrics can be found in the methods.

## Methods

### Study site

Our study utilizes data collected at the BEF-China experimental platform, a large-scale forest biodiversity experiment in subtropical China (Bruelheide *et al*. 2014). The experiment is located in Xingangshan, Jiangxi Province (29°05’00’’–29°07’43’’N,117°54’19’’–117°55’53’’E) and was established in 2009. The climatic conditions at the experiment are characterized by a mean annual temperature of 16.7 °C and a mean annual precipitation of 1800 mm (Yang *et al*. 2013). Diverse subtropical forests are the natural vegetation at the two experimental sites. The experiment was planted in 2009 (site A) and 2010 (site B) using a species pool of 40 locally occurring tree species. The experiment comprises a total of 566 plots, each characterized with a specific tree composition that follow a diversity gradient ranging from monocultures to 24-species mixtures. Trees were planted on a 20 x 20 grid, for a total of 400 trees per plot. Tree species were randomly assigned to planting positions. Each plot covers an area of 25.8 x 25.8 m².

### Capturing tree species’ functional niches

#### Collecting and processing functional trait data

To characterize the functional niche of tree species, we primarily utilized leaf functional trait data collected in 2018 (site A) and 2019 (site B). A total of 284 trees were sampled. Trees were selected to represent the entire range of environmental conditions within the experiment, and in particular the diversity treatment. Leaf traits were collected at varying heights of the tree crown to account for within-tree trait variation, which can account for about 25% of trait variation within species on average (Davrinche *et al*. 2023). The functional leaf traits collected were chosen to represent the leaf economic spectrum (Wright *et al*. 2004), comprising structural and nutritional leaf traits. Specifically, we included leaf dry matter content (LDMC), specific leaf area (SLA), carbon to nitrogen ration (C:N), as well as phosphorus (P), potassium (K), magnesium (MG), and calcium (CA). Detailed sampling and data processing protocols can be found in (Davrinche & Haider 2021).

To guide interpretations of major trait gradients, we supplemented the traits collected in 2018/19 with additional trait data collected in 2014 that cover an overlapping but broader set of leaf functional traits, but followed a sampling protocol that does not include within-individual variation due to being sampled from the bottom of the tree crown. Despite slightly different sampling schemes, the measurement of traits followed very similar protocols and are therefore comparable. In addition to the traits already included within the data from 2018/19, the data from 2014 also include measures of cellulose, hemicellulose, lignin, phenol, and tannin concentrations in the leaf tissue.

To capture tree species functional niches, we used the trait data collected in 2018/19 in a principal component analysis (PCA). All traits were standardized, and LDMC and C:N were log-transformed before calculating the PCA. To fill minor gaps in the data (1.3%), we imputed missing trait values using an approach based on random forest models (Stekhoven & Bühlmann 2012). To capture within-species variation, we used trait measures from multiple trees per species as our sample unit. For our analysis, we focused on the first two principal component axes, which had a larger eigenvalue than the mean eigenvalue of all principal component axes (Kaiser-Guttman Criterion). Together, these two axes have a cumulative eigenvalue of >80%.

Utilizing the results of the PCA, we then calculated six indices representing different aspects of the functional niche. First and second, we recorded the projection of species centroids on the two principal components to capture functional identities of species. To interpret these values, we compared the species positions to the loadings of the functional traits (see Fig. 1a). To guide interpretations, we fitted additional traits using data collected in 2014. Third, we recorded the absolute distance of a species’ centroid to the centre of gravity of trait space to measure functional specialization. This allows us to assess how extreme or common a species’ ecological strategy is compared to the average species (Bellwood *et al*. 2006). Fourth, we measured within-species variation as functional richness, which quantifies the convex hull volume of a species that is spanned up by the individual trees (Villéger *et al*. 2008). Fifth, we quantified if species utilize their trait space evenly or if specific strategies dominate it by calculating functional evenness (Villéger *et al*. 2008). Finally, we measured functional uniqueness of a species by determining the maximum relative overlap of convex hulls with other species, and subtracting the overlapping proportion from one (adapted from Villéger *et al*. 2011). Together, these indices allow us to quantify a species’ functional niche by quantifying its identity (position along principal component axis, functional specialization), variability (functional richness, functional evenness) and (dis)similarity with other species (functional uniqueness).

Because the number of sampled tree individuals varied slightly between species, we decided to standardize sampling efforts by resampling all species to a total of 10 individuals per species. We resampled 1000 times. To assure comparability, we resampled tree individuals from the PCA calculated over all tree individuals. Functional niche indices were calculated for each of the resampling runs, and then averaged.

### Assessing tree species’ interaction roles

#### Constructing species interaction networks

Our study comprises three types of interaction data, with interacting species groups that utilize the tree leaves as main resource and habitat. Naturally, sampling protocols differed between interaction types, but were standardized within interaction type and plots, allowing direct comparisons. The three interaction types include leaf chewing Lepidoptera larvae, leaf sucking Hemiptera, and endophytic bacteria. Interaction data was sampled across the entire diversity gradient.

Lepidoptera larvae were sampled in 2018 on 51 plots on both experimental sites (Wang *et al*. 2019). In each sampled plot, 80 trees were beaten with a padded stick, and falling Lepidoptera larvae were collected on a white sheet. Lepidoptera larvae were identified using meta-barcoding. Hemiptera sampling was carried out on 303 plots at both experimental sites (Fornoff *et al*. 2019). Species were visually identified directly on the leaves in 2014. The number of trees sampled per plot differed between diversity treatments, with a minimum of 36 trees being assessed in monocultures. Finally, endophytic leaf bacteria were sampled in 2019 from 10-20 shade leaves collected from a minimum of 8 trees per plot and across 29 plots (Yang *et al*. 2023). All sampling plots were located on site A. Endophytic bacteria were identified using meta-barcoding. For detailed sampling protocols of the three interaction types, see Wang *et al*. (2019), Fornoff *et al*. (2019), and Yang *et al*. (2023).

To analyse the interaction role of tree species, we constructed species interaction networks from the available interaction data. We constructed one interaction network per interaction type, site, and diversity level. This allowed us to explicitly integrate tree species richness in our analyses, and account for variation between sites and interaction types. Because sampling efforts varied between sites, we decided to standardize the numbers of trees sampled per species and network using a resampling approach. Due to differences in the sampling protocols and the densities of the different taxa, the number of trees differed between interaction types, with Lepidoptera using 4, Hemiptera using 80, and endophytic bacteria using 2 trees per species and network. For each network, we resampled 1000 times and calculated indices of interaction roles each time. We then averaged the indices for our analyses to gain a single value per species and network.

#### Indices capturing species’ interaction roles

For each network, we calculated six species-based indices that capture different aspects of species interaction roles. First, we quantified a tree species’ degree as its number of interaction partners, capturing the variability of a tree species interaction role. Second, we calculated the species’ identity using a weighted closeness measure (Dormann *et al*. 2009). Using a network projection, closeness quantifies how close a tree species is to the other tree species in the network. Weights were assigned based on abundance differences of interaction partners. Third, we calculated betweenness using a similar approach, but instead of focusing on how close a tree species is to other tree species, we measured how often the focal tree species is on the shortest path between all pairs of the other tree species. Fourth, we calculated interaction evenness of tree species as Shannon evenness of its weighted interactions, capturing variability of tree species interaction roles. Fifth, similarities of interactions were calculated by measuring the similarity of interactions between species pairs based on Morisita-Horn dissimilarities (Horn 1966). A species value was then calculated by taking the average similarity with other species. Finally, we calculated consumer dependence as the average relative dependence of consumer species on the focal tree species, with the relative dependence being calculated as the weighted proportion of interactions with the focal tree species compared to all interactions of the consumer species (Albert *et al*. in prep). Consumer dependence also captures species similarities, but from the consumer perspective.

### Tree biomass

To assess tree species biomass, we calculated above ground wood volume [m³] from annual census data. The data was collected each year at the core of each plot, with the core area comprising the central 6 x 6 (monoculture and 2-species mixtures) and 12 x 12 (4-, 8-, 16-, and 24-species mixtures) tree individuals of a plot. For the census data, the basal area and height of each tree was measured, which we then multiplied together with a cylindrical form factor of 0.5 to calculate above ground wood volume (Fichtner *et al*. 2017). We matched the tree biomass data to the sampling years and plots of the interaction data, allowing us to calculate interaction type-specific tree biomasses for each tree species in their respective tree species richness levels and sites.

### Statistics

To test how functional niches affect interaction roles of tree species, we build two models for each of the six response variables describing the different aspects of tree species’ interaction roles. The first model focuses on the effect of species identity captured as the position along the principal component axes, whereas the second model focuses on the remaining four variables. We split the analyses in two to avoid overfitting and account for intrinsic correlations between functional specialization and the functional identity based on PC axes. Each model initially contained the functional niche metrics in interaction with tree species richness and tree biomass, as well as crossed random effects accounting for differences between sites, network type, and tree species. All models were fitted in a linear modelling framework using the glmmTMB package (Brooks *et al*. 2017). We made sure linear assumptions were met using the performance package (Lüdecke *et al*. 2021). To make this possible, some variables had to be transformed. Specifically, consumer dependence, functional richness, tree species richness, and tree biomass were log-transformed prior to analyses. We furthermore used a Lambert W transformation to account for skewness and heavy tails in betweenness and interaction evenness (Goerg 2011). All variables were normalized before the analysis, with interaction metrics being normalized within interaction types to account for differences in scale.

## Results

We tested how functional niches of tree species determine their centrality in species interaction networks and found consistent responses across different centrality metrics (Fig. 2). Specifically, we found effects of the functional identity of tree species on centrality (Fig. 2, first column), indicating that species with more conservative growth strategies (i.e. higher values of the first principal component; Fig. 1a) tend to be more central in interaction networks. Network centrality also increased with the functional uniqueness of tree species (Fig. 2, second column), especially at high tree species richness. At low species richness, network centrality was independent and, for network closeness and betweenness, even negatively related to functional uniqueness.

**Fig. 2:**
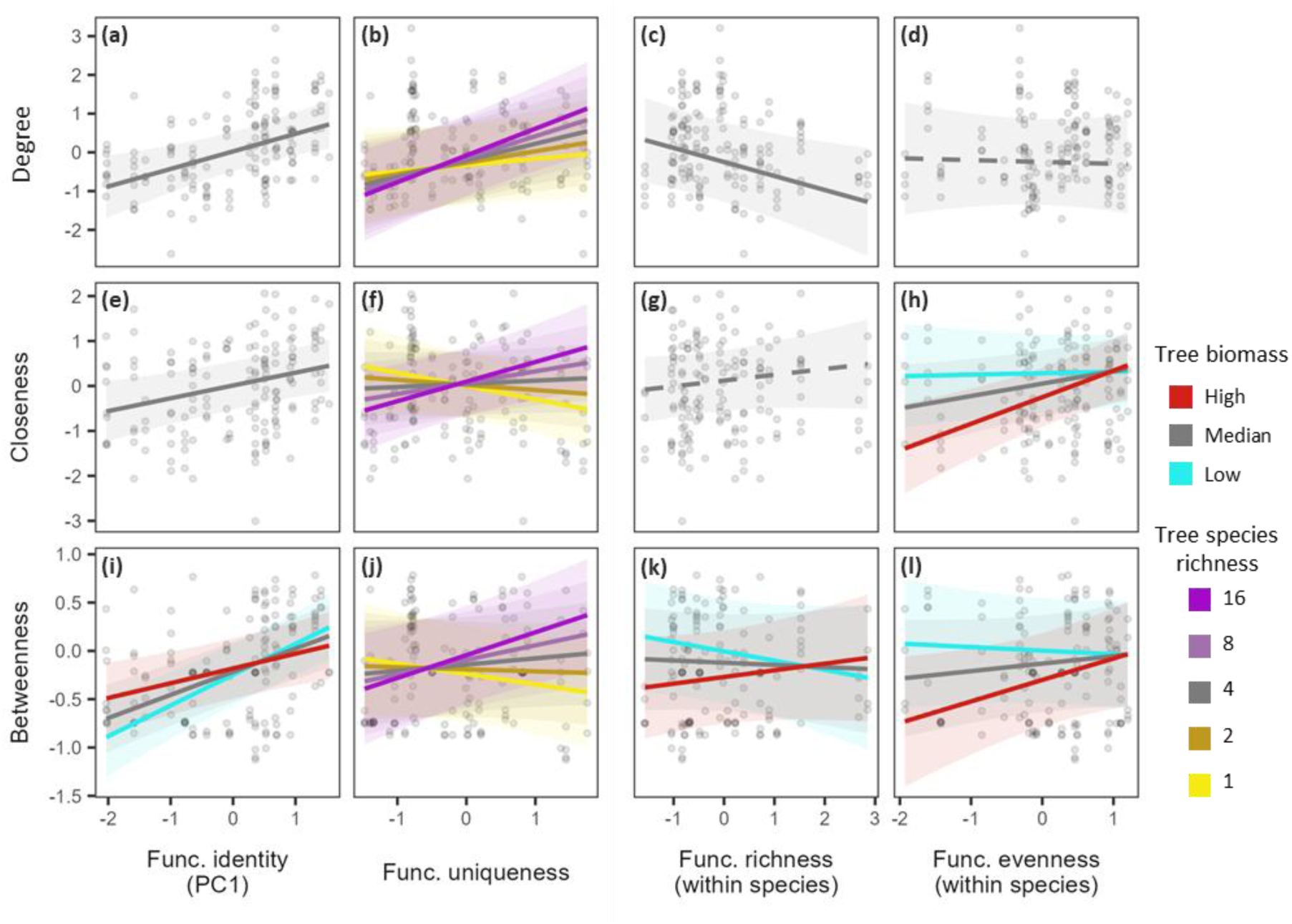
Effects of functional niche metrics on indices of network centrality of tree species. Effects are shown for the three centrality measures (degree, closeness, betweenness), and selected functional niche metrics that had effects on more than one centrality measure. Functional niche metrics comprise functional identity based on the first (PC1) principal component axis, functional uniqueness, functional richness, and functional evenness. Functional richness and evenness are based on multiple measures per species, thus capture within-species variability. Coloured lines show interaction effects with tree species richness and tree biomass. Solid lines indicate significant effects and dashed lines indicate non-significant effects (threshold: p = 0.05). See Tables S2 & S3 for model statistics.

Interestingly, closeness and betweenness, capturing the relative position of tree species in species interaction networks, and degree, capturing the variability of interactions, also showed differentiated response to differences in functional niches (Fig. 2, third and fourth column). While the functional richness of a species, capturing within-species variation, clearly reduced degree (Fig. 2c), such effects were not significant for closeness (Fig. 2g) and only emerged for small tree species for betweenness. For more productive species, effects ranged from neutral to positive (Fig. 2k). Similarly, only degree showed a response to functional specialization, indicating that species with more extreme ecological strategies, especially when highly productive, interact with fewer species (Tab. S2). In contrast, functional evenness, which captures how evenly tree individuals are distributed within the functional space occupied by a species, was not significant for degree (Fig. 2d), but showed very similar patterns for closeness and betweenness. The interaction effect reveals that, especially among highly productive species, functional evenness increases network centrality (Fig. 2h, l), an effect that was only neutral for the least productive species.

With the exception of functional specialization, all tested indices of tree functional niches showed effects on similarities of interactions between tree species, often showing interaction effects with tree biomass, tree species richness, or both (Fig. 3). Whenever we found interaction effects, average trends of functional niches were relatively weak (grey lines in Fig. 3a-c, e, f), but more pronounced effects occurred at the extremes of tree biomass and tree species richness. Effects of tree species’ growth strategies (PC1, Fig. 1a) show that large trees with conservative growth strategies tend to have more similar interactions, whereas smaller trees with conservative growth strategies tend to have less similar interactions (Fig. 3a). Differences in the composition of structural leaf components (PC2, Fig. 1a) had the strongest effects on large trees and trees growing in monoculture, where species with tougher leaves (high lignin, low hemicellulose) had more similar interactions than species with softer leaves (Fig. 3b, e). Tree species with evenly distributed functional strategies (higher functional evenness) showed a higher similarity of their interactions, especially for large trees and trees growing in monocultures. Interestingly, the effects of tree species richness and tree biomass seem to counteract each other for functional identity (PC2, Fig. 3b, e) and functional evenness (Fig. 3c, f), a trend also observed in interaction with effects of functional uniqueness on the degree of tree species (Tab. S3). Independent from tree biomass and tree species richness, functional richness, capturing tree species’ intraspecific variability, and functional uniqueness, capturing the uniqueness of a species’ functional niche compared to the other tree species, showed the clearest trends. While functional richness increased the similarity of interactions between tree species (Fig. 3d), functional uniqueness reduced it (Fig. 3g).

**Fig. 3:**
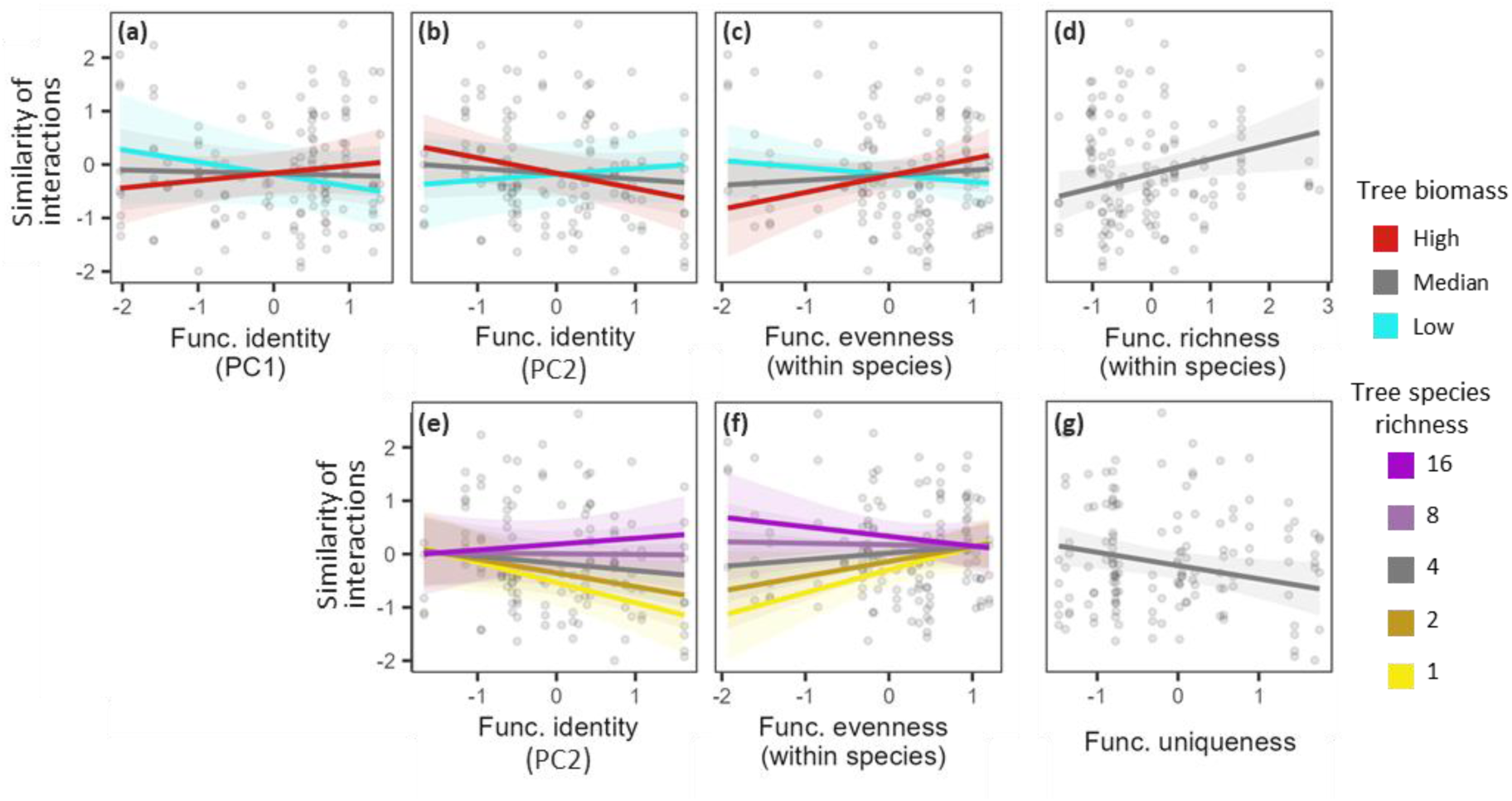
Similarity of interactions between tree species responds to changes in functional niches, tree species richness, and tree biomass. All significant (p < 0.05) effects of functional niche metrics are shown, i.e. the functional identity based on the first (PC1) and second (PC2) principal component axis, functional evenness, functional richness, and functional uniqueness. Functional richness and evenness are based on multiple measures per species, thus capture within species variability. Coloured lines show interaction effects with tree species richness and tree biomass. Interactions with tree biomass are shown in top row, interactions with tree species richness in bottom row. See Tables S2 & S3 for model statistics.

Not just the interactions of tree species, but also the dependence of consumers on the tree species changed with tree functional niches. For small tree species, the functional identity, specifically the structural composition of the leaves, had the clearest effects, whereas large trees showed no trends but overall a higher mean consumer dependence (Fig. 4a, Fig. 1a). In contrast, functional evenness of small trees had no effects on consumer dependence, whereas consumers depended most on large trees when functional evenness was low (Fig. 4b). Functional uniqueness did not affect consumer dependence at low tree species richness, but consumers of more functionally unique tree species were less dependent at higher diversity (Fig. 4c).

**Fig. 4:**
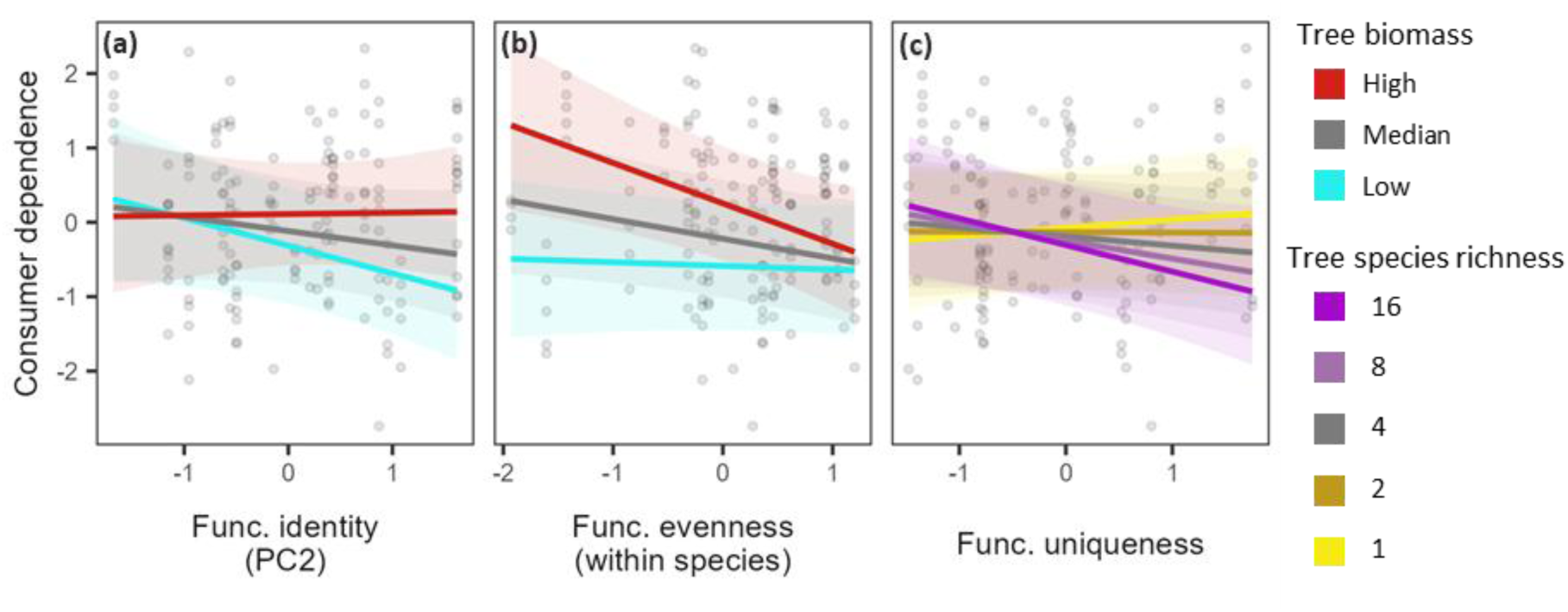
Effects of functional niche metrics on tree species’ consumer dependence. All significant (p < 0.05) effects of functional niche metrics are shown, i.e. the functional identity based on second (PC2) principal component axis, functional evenness, and functional uniqueness. Functional evenness is based on multiple measures per species, thus capture within species variability. Coloured lines show interaction effects with tree species richness and tree biomass. See Tables S2 & S3 for model statistics.

Interestingly, we found that interaction evenness of tree species showed no response to functional niches. However, it increased with tree biomass (Tab. S2 & S3), indicating that larger tree species tend to interact more evenly with their interaction partners.

## Discussion

To test how functional niches of tree species affect species’ interaction roles, we integrated species interaction data of three leaf-associated interaction types and leaf functional trait data in a joint analytical framework. Our analyses revealed that functional niches consistently alter interaction roles, particularly affecting network centrality and similarities of interactions. While the effects of functional niches on interaction roles often included direct associations between the same aspects (identity, variability, and similarities; H1), they were not limited to these direct associations (H2). Instead, functional niches showed a wide range of effects that often also depended on tree species richness and tree biomass (H3). While tree biomass had interactive effects in a wider range of relationships, effects of tree species richness were consistently related to metrics capturing species similarities. Our findings therefore demonstrate that the tight bond between tree species’ functional niches and the role they play in species interaction networks is also moderated by species’ biomass and may underpin niche differentiation processes that change with tree species richness.

### Similar interactions are determined by multiple aspects of functional niches

We found several matching aspects of functional niches and interaction roles of tree species, but the associations were less strict than expected. Most noticeably, functionally unique tree species had more unique (less similar) interactions compared to functionally more similar tree species. A tree species’ ability to avoid competition and contribute uniquely to an ecosystem therefore appears coordinated across functional niche and interaction role. This should naturally affect niche partitioning, the mechanism often ascribed to the usually positive biodiversity-ecosystem functioning relationships observed in forests (Liu *et al*. 2026). However, this may ultimately have limited impact, as niches do not need to be partitioned for a system to be diverse and highly functional (Albert *et al*. 2026), or may only need to be partitioned in limiting but not all dimension (Tilman 1982). Furthermore, competition in interaction networks is determined not just by plants but across interacting species groups that can fundamentally alter the outcome of competition (Albert *et al*. 2022; Brose 2008). Correspondingly, we found that consumer dependence did not simply follow changes in similarities of interactions of tree species, but instead showed an opposite trend, especially when tree species richness was high. While tree species with more unique traits had less unique interaction partners, interaction partners became less dependent on the tree species. This may be explained by the fact that functionally unique species had not only more unique, but generally a higher number of interactions, especially at high tree species richness. The fact that more diverse tree communities tend to enhance diversity throughout the ecosystem likely plays a role in this (Albert *et al*. 2026; Li *et al*. 2024). Despite an apparent link of tree species’ competitive abilities between functional niches and interaction networks, our findings thus suggest that the consequence of these links for species performances and biodiversity effects may differ drastically and need to be understood in the larger community context.

Similarities in tree species’ interactions responded not only to functional uniqueness but to most aspects of the functional niche. This further supports the idea that tree species interactions are especially sensitive to changes in the functional niche when focusing on the comparison between species (i.e. in the community context). Specifically, we found that species with more variable ecological strategies that occupy a broader functional niche space (higher functional richness) tend to share their interaction partners with other species. It could be expected that this effect is rooted in a simple geometrical association between niche size and similarity. However, we found that tree species with more variable functional niches (higher functional richness) actually interact with fewer species (lower degree). This effectively limits the space a species occupies in interaction networks, thus rendering the geometrical association less likely. Instead, species with large functional niches attract fewer interaction partners, possibly because a less focused functional strategy is less attractive as resource or habitat, especially for specialized interaction partners. The variability of functional niches could therefore help avoid negative interactions that can be additionally diluted as they are shared with other species (Root 1973). This, however, comes at the cost of potentially stronger competition due to having more similar interactions.

Interestingly, similarities in interactions, and thus the competitive ability of tree species, was most responsive to differences in functional niches in low diversity stands, where inter-specific competitive abilities matter the least. The reduced interaction similarity of tree species with a dominant functional strategy (low functional evenness) we observed in low diversity stands is likely indicative of tree species having more specialized interaction partners. In contrast, a reduced interaction similarity of species that invest in softer leaves (higher values of PC2, with less lignin, more hemicellulose) when grown in less diverse forests is surprising, but may result from specialist species that prefer trees with softer and thus more accessible leaves. Additionally, higher species densities at lower species richness are also generally more attractive to more specialized interaction partners (resource concentration hypothesis; Root 1973). For both, functional evenness and investment in leaf structural components, the low similarity of interactions seems to vanish in more diverse forests, with spill-overs of interaction partners from different tree species in the neighbourhood likely playing a role in either case (Borremans *et al*. 2019). The comparably weak effects of functional evenness and the investment in leaf structure at higher tree species richness renders the involvement of either functional niche aspect in differentiation processes of species interactions less likely. In contrast, the functional uniqueness of species tends to have the most pronounced effects on almost all tested aspects of tree species’ interaction roles, specifically in more diverse forest stands. It is therefore possible that the differentiation of tree species due to competition in diverse forest stands is largely realized in functional space, often more associated with how species take up and utilize resources, especially when based on leaf traits as in our study (Wright *et al*. 2004). Differentiation process of tree species interaction roles, where trees provide habitat and resources, seem in contrast less relevant for biodiversity effects, and may even inverse the often positive effects of biodiversity by limiting ecosystem functioning (Albert *et al*. 2026).

### Consistent effects across measures of network centrality

In addition to similarities, the positions of tree species in functional niche space and interaction networks showed signs of being coordinated as well. In particular, the tree species’ position in an interaction network, captured by the network centrality measures closeness and betweenness, changed with the species’ growth strategy, i.e. its position in functional trait space. But functional specialization, which captures how extreme an ecological strategy of a tree species is and thus resembles a centrality definition more closely, did not show any significant effects on the network centrality measures. At the same time, degree, which is closely related to the other two centrality measures (Jordán *et al*. 2007) but should be better at capturing the variability of interactions than the position in interaction networks, responded very similarly to changes in niche position. The similarities across centrality measures, however, are not consistent. Instead, tree species with more variable functional strategies (higher functional richness) have less variable (i.e. fewer) interactions, a pattern that is not reflected in closeness and betweenness. The observed effect that species with smaller functional niches tend to interact with more species is surprising in itself, and indicates that translating functional niches into interaction patterns might require a deeper understanding of the concrete processes that link them. Our initial expectation that tree species occupying more niche space are also attractive to more species relied on the assumption that interaction partners are largely generalists, a pattern found for consumer species in our experiment (Zhang *et al*. 2017). Our findings, however, indicate that specialist species may play a more impactful role in defining a tree species’ interaction role.

Despite some differences between degree and the other two centrality measures, effects of functional niches on network centrality were often strikingly similar across metrics. Our findings show that the centrality of tree species in interaction networks was higher when trees follow more conservative growth strategies. Increased herbivory rates on conservative trees have been previously observed in our study system (Schuldt *et al*. 2017) and may thus be a direct consequence of the high centrality of the species. But not just the tree growth strategy influenced all centrality metrics similarly, also the functional uniqueness of the tree species did. Average effects were positive, especially for degree, aligning with previous findings in pollen transport networks (Coux *et al*. 2016). However, functional uniqueness had neutral and even slightly negative effects on the network centrality of tree species when tree species richness was low. In contrast, at high tree diversity, functionally unique species were particularly central in the network. The higher centrality of both, functional unique tree species in diverse forests and tree species with conservative growth strategies, effectively renders them as connector species that are known to mediate environmental changes and disturbances (Martins *et al*. 2024). Given the consistency across centrality measures, effects of functional niches may therefore be suitable to identify species that critically determine ecosystem stability.

### Tree biomass moderates functional niche effects

Our findings revealed that many effects of functional niches on interaction roles of tree species were moderated by tree biomass. We anticipated effects of tree biomass because more productive, larger trees are likely attracting more interactions simply by being bigger (i.e. sampling effects), which would result in gradual shifts in the strength of the same trends. However, more productive tree species did not seem to attract more interaction partners, as effects of tree biomass on degree did not display any clear trends. Instead, tree biomass gradually shifted trends in interaction with functional evenness. Specifically, less productive species showed only weak and even neutral effects on multiple aspects of species interaction roles. In contrast, interaction roles of high-biomass tree species were particularly distinct at low functional evenness, indicating that tree species with dominant functional strategies (low functional evenness) have more distinct interaction roles. Specifically, high-biomass tree species with dominant functional strategies were less central in interaction network (low closeness and betweenness), had less similar interactions, and their consumers depended more on them. Together, these findings indicate that tree species with large biomass and dominant functional strategies attract more distinct, more specialized interaction partners. A focused functional strategy can be the result of a favourable ecological strategy, that, for example, allows a better defence against herbivory. That species with more focused functional strategies attract more specialized consumers is therefore not necessarily surprising. The fact that such effects do not emerge for less productive species is, however, noteworthy, as it indicates that the potential benefits of a focused strategy can only emerge when a tree species is already growing well. In turn, this implies that species that grow less optimally, for example due to being less well adapted to environmental conditions, have also less room to optimize interactions.

Unless when interacting with functional evenness, our findings show that effects of tree biomass are often more differentiated, with more productive trees showing opposite trends to less productive trees. Such contrasting effects imply that multiple overlapping processes shape tree species responses. Identifying these processes is, however, difficult because biomass can act as a proxy for other effects. While our analytical approach aimed at controlling environmental heterogeneity by aggregating networks at the site-scale, we cannot fully rule out that environmental drivers play a role in differentiating tree biomass effects. Tree biomass may also mask tree species richness effects, which enhances tree biomass in our experiment and in tree biodiversity experiments globally (Huang *et al*. 2018; Liu *et al*. 2026). By focusing on individual tree biomass, we could minimize tree density effects that critically influence diversity effects on stand productivity (Liu *et al*. 2026). Indeed, whenever we found simultaneous tree biomass and tree species richness effects, they were often inversed, with low biomass and high species richness (and vice versa) showing similar trends. Tree biomass and tree species richness therefore seem to capture different processes that indicate potential trade-offs defining a tree species interaction role. Ultimately, our findings show that the way tree biomass shapes how the functional identity of tree species shapes their interaction roles is much less clear cut than could be expected.

## Conclusion

Ecologists have long realized the importance of functional traits and, consequently, functional niches for determining species interactions (Brose *et al*. 2019; Peralta *et al*. 2024). We expanded on this idea by investigating how different aspects of functional niches are altering aspect of interaction roles of tree species in interaction networks of leaf-associated consumer species. Our findings clearly show that there are direct links within the three aspects – identity, variability, and similarity – between functional niche space and in interaction networks. However, a direct translation still seems difficult. Ecological drivers that determine tree species’ functional niche can differ from drivers determining interaction roles. We have, for example, discussed niche differentiation processes that seem to be more pronounced for functional niches than for interaction roles. This highlights the importance of conserving a network perspective, which finds further support in the fact that most metrics that describe the interaction role of tree species, and thus define their role in ecosystems (Albert *et al*. 2026; Cirtwill *et al*. 2018), showed simultaneous responses to multiple aspects of functional niches. Interactive effects with tree biomass and tree species richness furthermore highlight the sensitivity of interaction roles to the larger ecological context. Even though the effects of species with specific interaction roles on ecosystem dynamics is still not fully known, evidence of their benefits (e.g. network centrality for ecosystem stability; Martín González *et al*. 2010; Martins *et al*. 2024) emphasizes the potential of identifying drivers of interaction roles for anticipating ecosystem change.

## Acknowledgements

GA, MS, AS, HB, FF were supported by the MultiTroph Research Unit funded by the German Research Foundation (DFG, 452861007/FOR 5281). HB, AS, AD, SH, WH, GO, YL, XL, CZ acknowledge the support from the International Research Training Group TreeDì, jointly funded by the DFG (319936945/GRK2324) and the University of Chinese Academy of Sciences (UCAS). GA was additionally supported by the Chinese Academy of Sciences (CAS) President’s International Fellowship Initiative (2026PVC0132). We are grateful for support from the BEF-China platform, which was established with funds from the Deutsche Forschungsgemeinschaft (DFG, German Research Foundation; grant DFG 35758305/FOR 891), the National Natural Science Foundation of China (NSFC 30710103907, 30930005, 31170457 and 31210103910), and the Swiss National Science Foundation (SNSF). We thank all local workers and assistants who participated in data collection.

## Author contributions

GA and AS developed the idea, analysed the data, and led the writing. AD contributed to conceptualizing the idea. MS, FF, AK, AD, SH, TP, WH, GO, LJ, YL, XL, MW, PW, XY, CZ provided data and supported data preparation. GA wrote the first draft of the manuscript and all authors contributed to revisions.

## Data availability

The data underlying the analyses presented in this study will be made available on a public repository after acceptance of the manuscript.

## Notes

### Competing Interest Statement

The authors have declared no competing interest.

