## Supplementary information for "Functional niche attributes shape species interaction roles in ecological networks"

**Supplementary information includes:**

Supplementary Tables 1-3

### Supplementary Tables

**Table S1: Tree species used in the study and abbreviations of species names as used in Fig. 1a.**

| Species name | Abbreviation |
|---|---|
| *Alniphyllum fortunei* | A.for |
| *Castanea henryi* | C.hen |
| *Castanopsis fargesii* | C.far |
| *Castanopsis sclerophylla* | C.scl |
| *Choerospondias axillaris* | C.axi |
| *Cinnamomum camphora* | C.cam |
| *Cyclobalanopsis glauca* | C.gla |
| *Elaeocarpus chinensis* | E.chi |
| *Liquidambar formosana* | L.for |
| *Lithocarpus glaber* | L.gla |
| *Machilus thunbergii* | M.thu |
| *Manglietia fordiana* | M.for |
| *Nyssa sinensis* | N.sin |
| *Quercus fabri* | Q.fab |
| *Quercus serrata* | Q.ser |
| *Rhus chinensis* | R.chi |
| *Sapindus saponaria* | S.sap |
| *Schima superba* | S.sup |
| *Triadica sebifera* | T.seb |

**Table S2: Model statistics of functional identity models.** Significant p values ($p < 0.05$) are highlighted in bold. Explained variance (marginal): degree: $R^2 = 0.231$, closeness: $R^2 = 0.146$, betweenness: $R^2 = 0.189$, interaction evenness: $R^2 = 0.045$, similarity of interactions: $R^2 = 0.181$, consumer dependence: $R^2 = 0.157$.

| | Estimate | Std. Error | z value | p value |
|---|---|---|---|---|
| *degree* | | | | |
| (Intercept) | 0.033 | 0.262 | 0.126 | 0.899 |
| F. identity (PC1) | 0.453 | 0.142 | 3.191 | **0.001** |
| Tree wood volume [$m^3$] | -0.159 | 0.198 | -0.801 | 0.423 |
| *closeness* | | | | |
| (Intercept) | 0.021 | 0.274 | 0.076 | 0.940 |
| F. identity (PC1) | 0.285 | 0.090 | 3.150 | **0.002** |
| F. identity (PC2) | -0.035 | 0.128 | -0.271 | 0.786 |
| Tree wood volume [$m^3$] | -0.109 | 0.138 | -0.793 | 0.428 |
| F. identity (PC2):Tree wood volume [$m^3$] | -0.264 | 0.107 | -2.459 | **0.014** |
| *betweenness* | | | | |
| (Intercept) | -0.213 | 0.158 | -1.349 | 0.177 |
| F. identity (PC1) | 0.233 | 0.040 | 5.908 | **3e-09** |
| Tree wood volume [$m^3$] | 0.054 | 0.046 | 1.163 | 0.245 |
| F. identity (PC1):Tree wood volume [$m^3$] | -0.138 | 0.059 | -2.350 | **0.019** |
| *interaction evenness* | | | | |
| (Intercept) | 0.148 | 0.134 | 1.100 | 0.271 |
| Tree wood volume [$m^3$] | 0.222 | 0.089 | 2.500 | **0.012** |
| *similarity of interactions* | | | | |
| (Intercept) | -0.065 | 0.220 | -0.294 | 0.769 |
| F. identity (PC1) | 0.022 | 0.127 | 0.175 | 0.861 |
| F. identity (PC2) | -0.136 | 0.134 | -1.015 | 0.310 |
| Tree species richness | 0.254 | 0.068 | 3.756 | **2e-04** |
| Tree wood volume [$m^3$] | 0.114 | 0.155 | 0.738 | 0.460 |
| F. identity (PC1):Tree wood volume [$m^3$] | 0.257 | 0.119 | 2.164 | **0.030** |
| F. identity (PC2):Tree species richness | 0.202 | 0.077 | 2.618 | **0.009** |
| F. identity (PC2):Tree wood volume [$m^3$] | -0.282 | 0.114 | -2.472 | **0.013** |
| *consumer dependence* | | | | |
| (Intercept) | -0.110 | 0.362 | -0.304 | 0.761 |
| F. identity (PC2) | -0.185 | 0.187 | -0.987 | 0.324 |
| Tree wood volume [$m^3$] | 0.337 | 0.171 | 1.972 | **0.049** |
| F. identity (PC2):Tree wood volume [$m^3$] | 0.309 | 0.111 | 2.790 | **0.005** |

**Table S3: Model statistics of functional niche models.** Significant p values ($p < 0.05$) are highlighted in bold. Explained variance (marginal): degree: $R^2 = 0.196$, closeness: $R^2 = 0.176$, betweenness: $R^2 = 0.141$, interaction evenness: $R^2 = 0.040$, similarity of interactions: $R^2 = 0.180$, consumer dependence: $R^2 = 0.188$.

| | Estimate | Std. Error | z value | p value |
|---|---|---|---|---|
| *degree* | | | | |
| (Intercept) | -0.227 | 0.525 | -0.431 | 0.666 |
| F. richness (within sp.) | -0.361 | 0.152 | -2.381 | **0.017** |
| F. specialization | -0.352 | 0.154 | -2.287 | **0.022** |
| F. uniqueness | 0.427 | 0.153 | 2.789 | **0.005** |
| Tree species richness | 0.087 | 0.056 | 1.562 | 0.118 |
| Tree wood volume [$m^3$] | -0.129 | 0.193 | -0.670 | 0.503 |
| F. specialization:Tree wood volume [$m^3$] | -0.288 | 0.114 | -2.527 | **0.012** |
| F. uniqueness:Tree species richness | 0.184 | 0.062 | 2.968 | **0.003** |
| F. uniqueness:Tree wood volume [$m^3$] | -0.243 | 0.114 | -2.137 | **0.033** |
| *closeness* | | | | |
| (Intercept) | 0.019 | 0.367 | 0.053 | 0.958 |
| F. evenness (within sp.) | 0.315 | 0.127 | 2.484 | **0.013** |
| F. uniqueness | 0.067 | 0.116 | 0.576 | 0.565 |
| Tree species richness | 0.035 | 0.074 | 0.465 | 0.642 |
| Tree wood volume [$m^3$] | -0.453 | 0.219 | -2.069 | **0.039** |
| F. evenness (within sp.):Tree wood volume [$m^3$] | 0.454 | 0.153 | 2.970 | **0.003** |
| F. uniqueness:Tree species richness | 0.255 | 0.082 | 3.108 | **0.002** |
| F. uniqueness:Tree wood volume [$m^3$] | -0.227 | 0.120 | -1.884 | *0.060* |
| *betweenness* | | | | |
| (Intercept) | -0.156 | 0.253 | -0.619 | 0.536 |
| F. richness (within sp.) | -0.007 | 0.070 | -0.096 | 0.923 |
| F. evenness (within sp.) | 0.103 | 0.087 | 1.184 | 0.236 |
| F. uniqueness | 0.060 | 0.070 | 0.857 | 0.391 |
| Tree species richness | 0.068 | 0.034 | 1.993 | **0.046** |
| Tree wood volume [$m^3$] | -0.236 | 0.102 | -2.315 | **0.021** |
| F. richness (within sp.):Tree wood volume [$m^3$] | 0.130 | 0.057 | 2.278 | **0.023** |
| F. evenness (within sp.):Tree wood volume [$m^3$] | 0.205 | 0.087 | 2.355 | **0.019** |
| F. uniqueness:Tree species richness | 0.120 | 0.037 | 3.263 | **0.001** |
| *interaction evenness* | | | | |
| (Intercept) | 0.155 | 0.134 | 1.162 | 0.245 |
| Tree wood volume [$m^3$] | 0.207 | 0.088 | 2.351 | **0.019** |

**Table S3** – continued

| | Estimate | Std. Error | z value | p value |
|---|---|---|---|---|
| *similarity of interactions* | | | | |
| (Intercept) | -0.222 | 0.124 | -1.785 | *0.074* |
| F. richness (within sp.) | 0.259 | 0.112 | 2.305 | **0.021** |
| F. evenness (within sp.) | 0.139 | 0.169 | 0.823 | 0.410 |
| F. specialization | -0.033 | 0.114 | -0.290 | 0.772 |
| F. uniqueness | -0.232 | 0.111 | -2.098 | **0.036** |
| Tree species richness | 0.256 | 0.068 | 3.753 | **2e-04** |
| Tree wood volume [m³] | 0.020 | 0.108 | 0.187 | 0.852 |
| F. evenness (within sp.):Tree species richness | -0.206 | 0.095 | -2.180 | **0.029** |
| F. evenness (within sp.):Tree wood volume [m³] | 0.373 | 0.149 | 2.503 | **0.012** |
| F. specialization:Tree species richness | -0.126 | 0.075 | -1.682 | *0.093* |
| *consumer dependence* | | | | |
| (Intercept) | -0.169 | 0.399 | -0.424 | 0.672 |
| F. evenness (within sp.) | -0.298 | 0.141 | -2.110 | **0.035** |
| F. uniqueness | -0.117 | 0.114 | -1.021 | 0.307 |
| Tree species richness | -0.080 | 0.071 | -1.133 | 0.257 |
| Tree wood volume [m³] | 0.684 | 0.186 | 3.684 | **2e-04** |
| F. evenness (within sp.):Tree wood volume [m³] | -0.402 | 0.160 | -2.506 | **0.012** |
| F. uniqueness:Tree species richness | -0.163 | 0.077 | -2.125 | **0.034** |
